# Natural Language Processing for Classification of Acute, Communicable Findings on Unstructured Head CT Reports: Comparison of Neural Network and Non-Neural Machine Learning Techniques

**DOI:** 10.1101/173310

**Authors:** Falgun H. Chokshi, Bonggun Shin, Timothy Lee, Andrew Lemmon, Sean Necessary, Jinho D. Choi

**Author notes:** Corresponding Author:* Falgun H. Chokshi, M.D., M.S., Department of Radiology and Imaging Sciences, Division of Neuroradiology, Emory University School of Medicine, 1364 Clifton Road NE, Atlanta, Georgia 30322, Twitter: @FalgunChokshiMD. Presented as an oral paper at the ASNR 2017 Annual Meeting (Long Beach, CA).

## Abstract

**Background and Purpose:** To evaluate the accuracy of non-neural and neural network models to classify five categories (classes) of acute and communicable findings on unstructured head computed tomography (CT) reports.

**Materials and Methods:** Three radiologists annotated 1,400 head CT reports for language indicating the presence or absence of acute communicable findings (hemorrhage, stroke, hydrocephalus, and mass effect). This set was used to train, develop, and evaluate a non-neural classifier, support vector machine (SVM), in comparisons to two neural network models using convolutional neural networks (CNN) and neural attention model (NAM) Inter-rater agreement was computed using kappa statistics. Accuracy, receiver operated curves, and area under the curve were calculated and tabulated. P-values < 0.05 was significant and 95% confidence intervals were computed.

**Results:** Radiologist agreement was 86-94% and Cohen’s kappa was 0.667-0.762 (substantial agreement). Accuracies of the CNN and NAM (range 0.90-0.94) were higher than SVM (range 0.88-0.92). NAM showed relatively equal accuracy with CNN for three classes, severity, mass effect, and hydrocephalus, higher accuracy for the acute bleed class, and lower accuracy for the acute stroke class. AUCs of all methods for all classes were above 0.92.

**Conclusions:**

1. Neural network models (CNN & NAM) generally had higher accuracies compared to the non-neural models (SVM) and have a range of accuracies that comparable to the inter-annotator agreement of three neuroradiologists.
2. The NAM method adds ability to hold the algorithm accountable for its classification via heat map generation, thereby adding an auditing feature to this neural network.

**Abbreviations:**

NLP: Natural Language Processing
CNN: Convolutional Neural Network
NAM: Neural Attention Model
HER: Electronic Health Record

## Introduction

The radiology report offers a major source of unstructured data that can be mined using natural language processing (NLP) and applied towards predictive models assessing outcomes such as length of stay, mortality, resource utilization, and cost-analysis. NLP encompasses a range of powerful data science and computational linguistics methods to process such large text-based data sets.^1^ An increasing body of literature has focused on the uses of various NLP techniques in radiology reports. In a recent systematic review, Pons et al. categorized 67 studies on the use of radiology NLP into discrete groups: 1) cohort building for epidemiologic studies, 2) quality assessment for radiology practice, and 3) clinical support services.^2^

Early rules-based NLP methods have been used to text mine radiology reports to evaluate outcomes such as head CT diagnostic yield in intensive care unit (ICU) patients,^3^ tumor information extraction for liver tumors,^4^ or determining brain tumor status via MRI reports.^5^ These rules based methods, however, were dependent on identifying specific words and phrases based on human references and annotations of training set reports^1, 2^ and some were beholden to domain-specific medical lexicons and ontologies.^6, 7^

Recent advances in machine learning based NLP techniques have shown promise in reliably classifying findings in unstructured radiology reports without the limitation of annotating specific words or phrases or beholden to simple rules-based NLP. Initial work in this space suggests that reports from specific modalities and body regions could be grouped together before embarking on the non-trivial task of developing machine learning-based NLP systems.^8^ Moreover, the performance of both non-neural and neural network models has yet to be demonstrated using radiology reports containing acute and communicable findings.

Therefore, this study aimed to compare the performance of both non-neural and neural network based NLP methods on the document-level extraction of acute and communicable findings in a sample of ICU head CT reports without linkage to any medical language ontologies. We compared the methods’ ability to classify 5 categories of findings that would be communicated to ordering clinical teams in routine radiology practice, per Joint Commission on Accreditation of Healthcare Organizations (JCAHO) guidelines.^9^

## Materials and Methods

### Study Design and Radiology Report Databases

Our institutional review board approved this HIPAA-compliant retrospective study and granted waiver of consent.

The annotated dataset used in this study was part of a larger set used in a study evaluating diagnostic yield of head CT in altered mental status amongst intensive care unit (ICU) patients.^3^ Briefly, we searched our institution’s clinical data warehouse for all consecutive, final radiology reports for non-contrast head CTs performed for altered mental status (*International Classification of Diseases*, 9^th^ edition code 780.97) in our ICUs for the date range of July 2011 to June 2013. We identified 2,486 consecutive non-contrast head CT exams’ reports.

Of these exams, the first 1400 consecutive head CT reports were annotated independently and adjudicated collectively by 3 radiologists (2 attendings and 1 neuroradiology fellow) and this set served as the reference database (“ground-truth”). As adapted from Chokshi et al,^3^ each radiology report was classified for 5 categories: 1) study severity, 2) acute intracranial bleed, 3) mass effect, 4) acute stroke and 5) hydrocephalus using a scale of 0 (normal) to 2 (new or worsening finding that would warrant a phone call to the ordering team). We then analyzed the inter-reader agreement and performed kappa statistics.

Additionally, to develop the neural network algorithms described below, an additional set of 80,000 continuous head CT reports was identified after a data warehouse search for emergency department (ED) head CTs performed between January 1, 2015 to December 1, 2016. These reports were intentionally not annotated and strictly served to improve the semantic NLP abilities of the neural network algorithms.

Next, to evaluate the performance of the three machine learning algorithms for classification of acute, communicable findings on the reports, all findings that were scored 0 or 1 were grouped together were grouped a negative for acute, communicable findings and those scored 2 as positive for acute communicable findings. This conversion to a binary outcome system allowed us to train the algorithms to be more accurate for clinically relevant findings.

### Machine Learning Algorithms

#### Non-Neural Model

We used the linear classifier, Support Vector Machines (SVM) as the strong baseline non-neural model to compare with the neural network models. A SVM identifies the strongest mathematical boundary between positive and negative examples in the training data.^10^ We used a Bag-of-Words (BOW) representation to feature engineer the SVM’s ability to find the maximum boundary between positive and negative data points for a given classifier.^11^ Since 5 classes of report findings were annotated, 5 distinct SVMs were developed, one for each class.

#### Neural Network Methods

Two neural network models were developed using Convolutional Neural Networks (CNN) and Neural Attention Model (NAM) where NAM gives another level of optimization to a CNN (both described below). To increase the robustness in accuracy of word semantics in the neural networks for radiology report text, the 80,000 un-annotated head CT reports were pre-tokenized and processed using Word2Vec.^12^ Word2Vec is open-source software that converts raw text into word vectors represented in Cartesian space. This allows contextual relationships between words and phrases to be geometrically evaluated and their strength can be quantified.

### Convolutional Neural Network (CNN)

We used a single layer CNN model for document classification. The CNN represented the text input as an input matrix, then as featured vectors, followed by dense vectors, and finally a prediction of output (classifier result, such as acute hemorrhage or not). Since 5 classes of report findings were annotated, 5 distinct CNNs were developed, one for each class.

### Neural Attention Model (NAM)

We selected a NAM as a comparison method because NAMs have the unique ability to show the attention of the input source from which they made their prediction or classification. ^13, 14^ For example, on a report annotated as positive for new intracranial hemorrhage, the CNN may simply say the report is positive, however a NAM can produce the same prediction and a heat map of all the words it found important to make that decision. This latter “rationalization” feature makes NAM models highly attractive for machine learning based NLP in radiology, when compared to conventional CNNs. See **Figure 2** for an example of a heat map.

**Figure 2.**
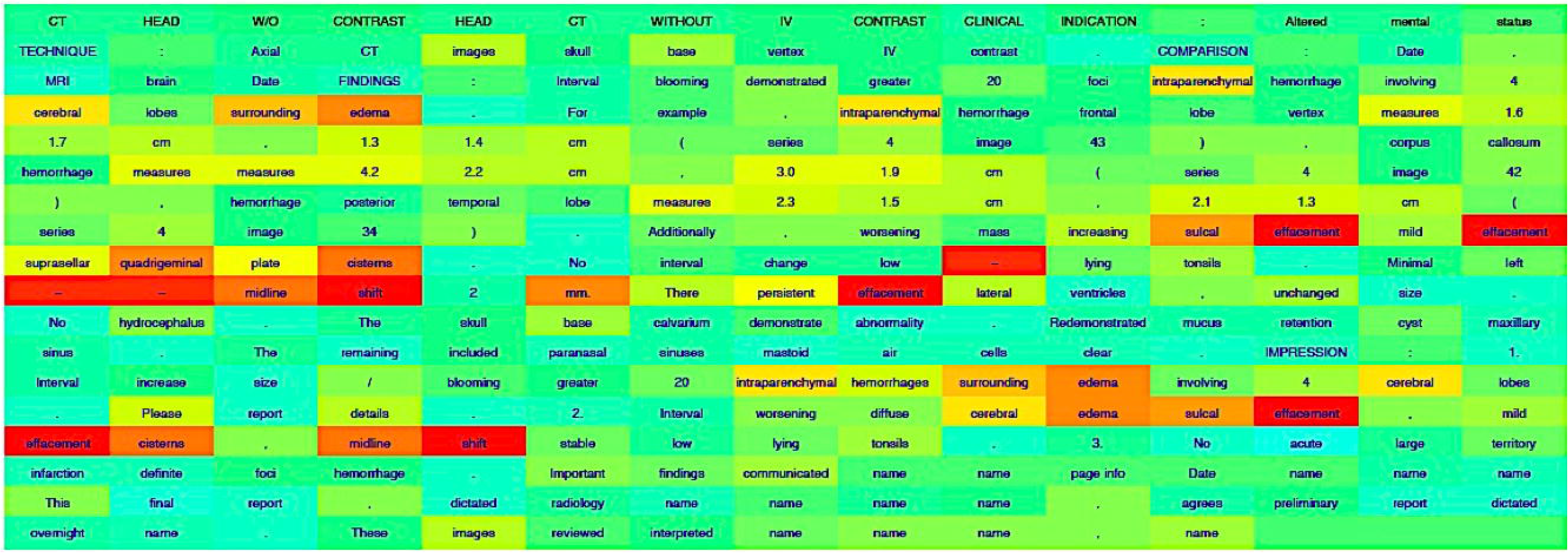
Heat Map of Head CT Report. Multi-color heat map generated from a single head CT report showing the terms used to make classification by the NAM in red. This report was classified as positive for mass effect. NAM, neural attention model.

The NAM architecture is an elaboration of the CNN model we used and involves an additional Attention Matrix layer and an Attention Vector layer imbedded in the CNN model at large (**Figure 1**).^13^ Similar to the CNN model, since 5 classes of report findings were annotated, 5 distinct NAMs were developed, one for each class.

**Figure 1.**
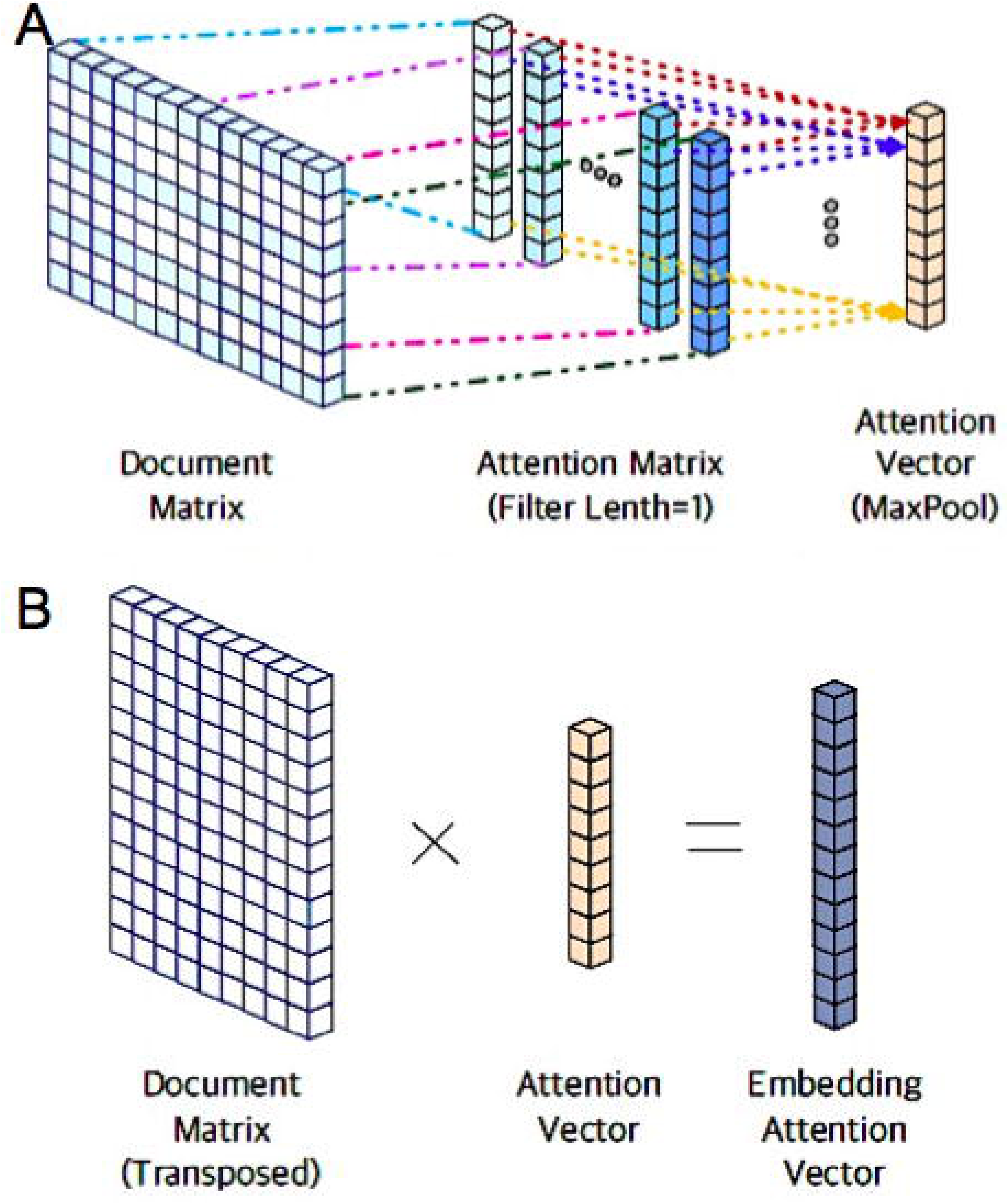
Neural Network Architectures. The single-layer CNN model is represented in (A) and is comprised of the document matrix, the attention matrix, and the attention vector. The single-layer NAM model is represented in (B) and is comprised of document matrix and attention vector, which combined form the embedding attention vector. CNN, convolutional neural network; NAM, neural attention model.

### Evaluation of Annotated Reports & Statistical Analysis

The annotated and adjudicated set of 1400 head CT reports was randomly divided into groups of 1000, 200, and 200, for training, validation, and testing sets, respectively. The same sets of 1000, 200, and 200 reports were used for the training, validation, and testing of the SVM, CNN, and NAM models for all 5 classes.

The primary metric was accuracy, which was measured by dividing agreed finding on annotation by the total finding on annotation. Using receiver operator curves (ROC) were calculated the area under the curve (AUC) for the three methods as well. Confidence intervals were determined at 95% and p-values <0.05 were significant.

## Results

### Radiologist Agreement

The three readers agreed 86-94% of the time and unweighted kappa scores (Cohen’s kappa) were between 0.667 and 0.762, showing substantial agreement.^15^

### Dataset Characteristics and NLP Metrics

**Table 1** shows the characteristics of the 5 classes in the annotated dataset of 1400 head CT reports. **Table 2** shows the accuracy of the three machine learning methods. **Figure 3** shows the ROC curves of the three methods and their associated AUC values.

**Table 1.** Characteristics of Acute Findings by Class for the Annotated Head CT Reports. 0, completely normal study; 1, abnormal findings but not acute and communicable; 2, abnormal findings that are acute and communicable.

| Class | Scores |  |  | Total |
| --- | --- | --- | --- | --- |
|  | 0 | 1 | 2 |  |
| **Severity of Study** | 58 | 940 | 402 | 1400 |
| **Acute Blood** | 653 | 546 | 201 | 1400 |
| **Mass Effect** | 751 | 443 | 206 | 1400 |
| **Acute Stroke** | 1113 | 173 | 114 | 1400 |
| **Hydrocephalus** | 1078 | 172 | 150 | 1400 |

**Table 2.** Performance Accuracy for Non-Neural and Neural-Network Machine Learning Models. Data are Percentage (95% Confidence Intervals). SVM, Support Vector Machine; CNN, Convolutional Neural Network; NAM, Neural Attention Model.

| Class | Accuracy |  |  |
| --- | --- | --- | --- |
|  | SVM | CNN | NAM |
| **Severity of Study** | 0.88 (0.83,0.92) | 0.91 (0.87,0.95) | 0.91 (0.87,0.95) |
| **Acute Blood** | 0.87 (0.82,0.91) | 0.93 (0.89,0.97) | 0.94 (0.90,0.97) |
| **Mass Effect** | 0.92 (0.88,0.96) | 0.90 (0.86,0.94) | 0.92 (0.88,0.96) |
| **Acute Stroke** | 0.91 (0.87,0.95) | 0.94 (0.91,0.97) | 0.93 (0.90,0.97) |
| **Hydrocephalus** | 0.91 (0.87,0.95) | 0.92 (0.88,0.96) | 0.93 (0.89,0.96) |

**Figure 3.**
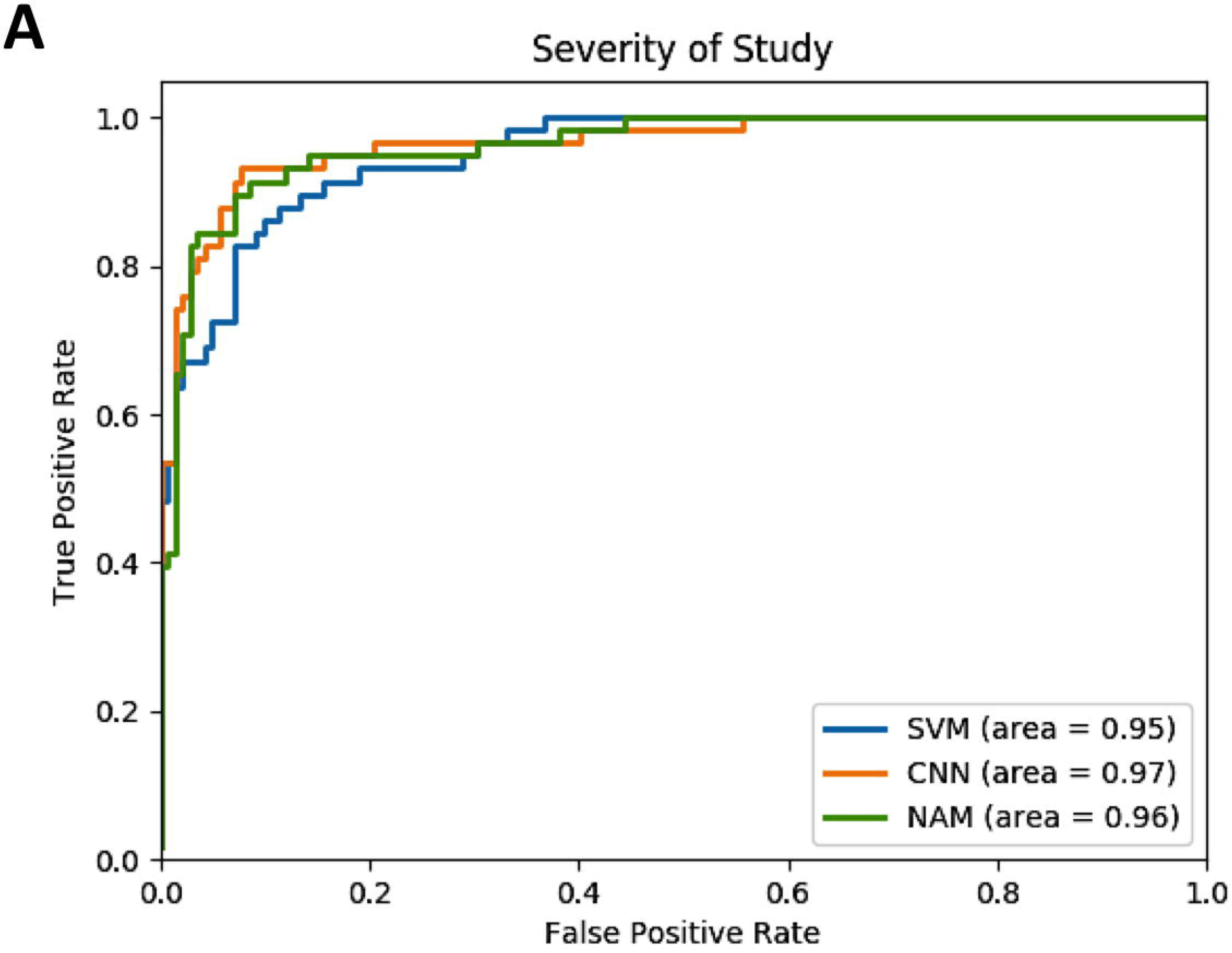
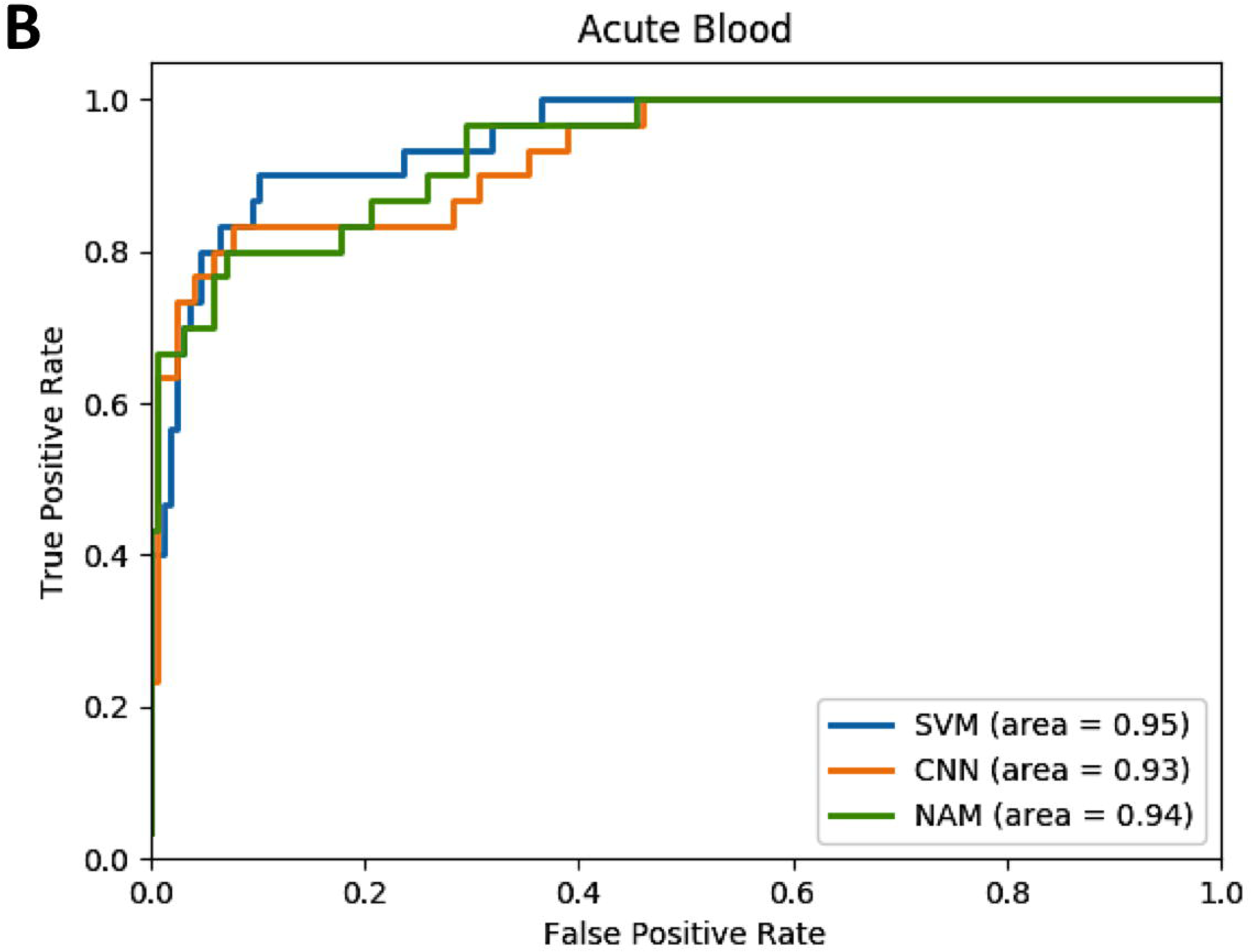
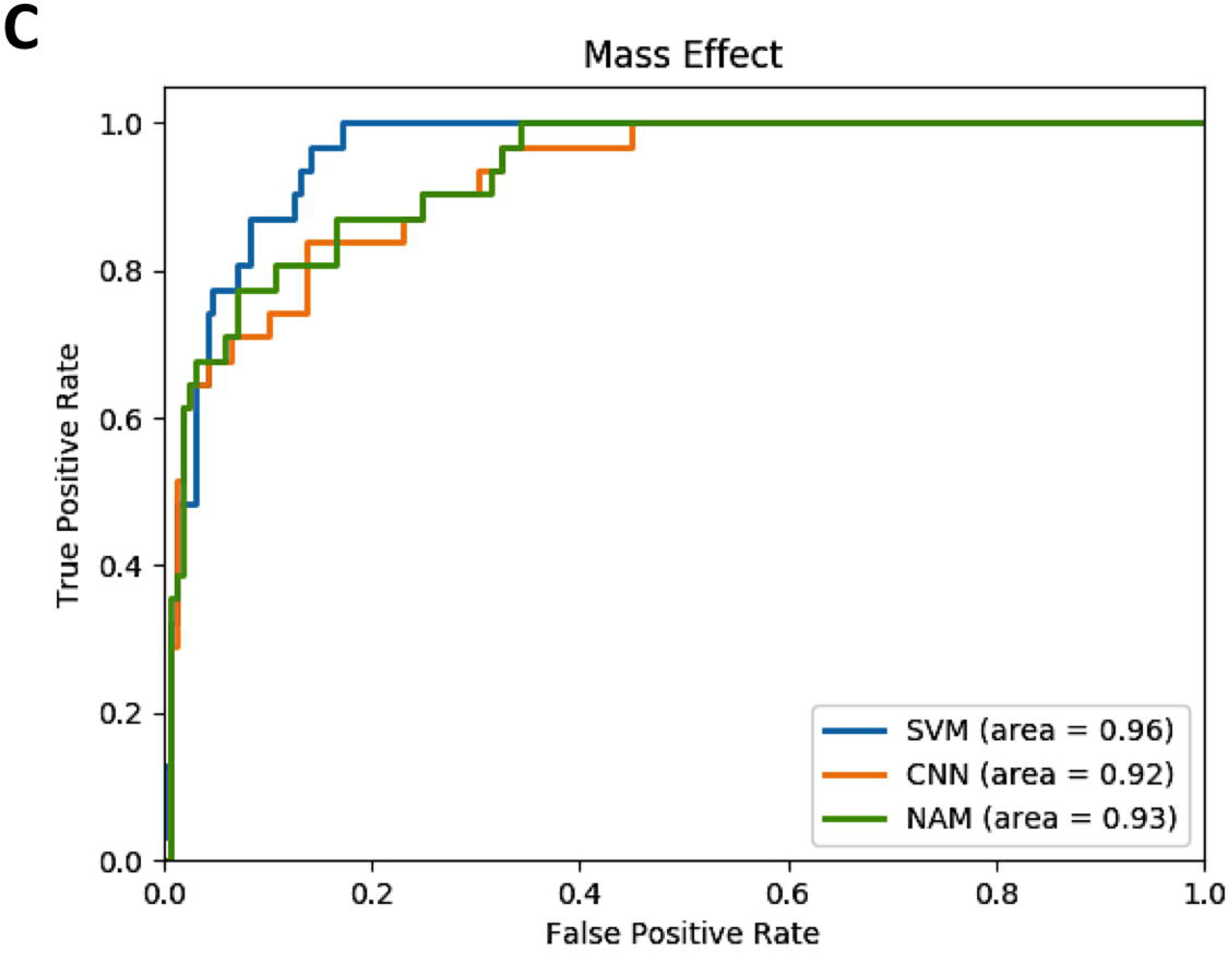
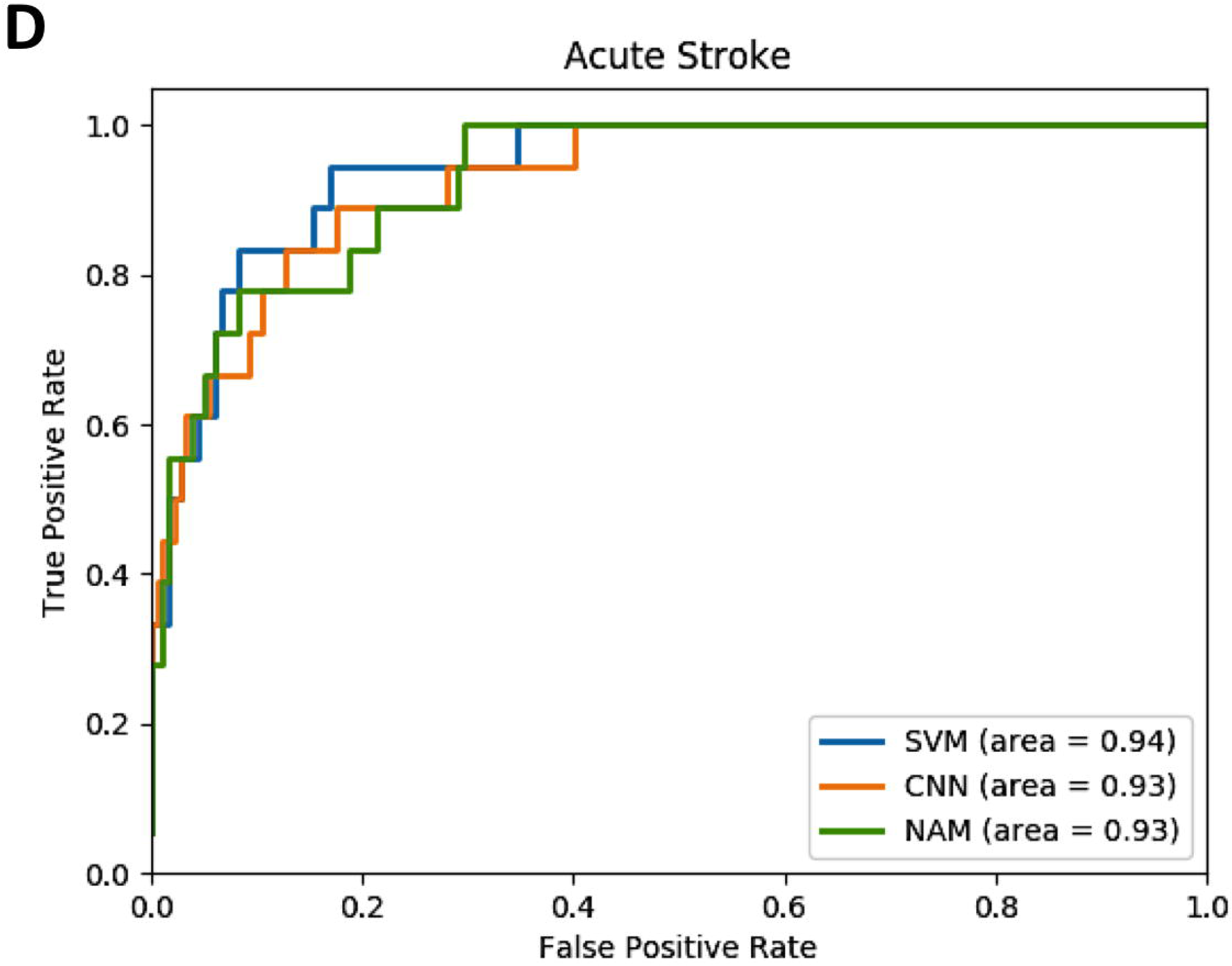
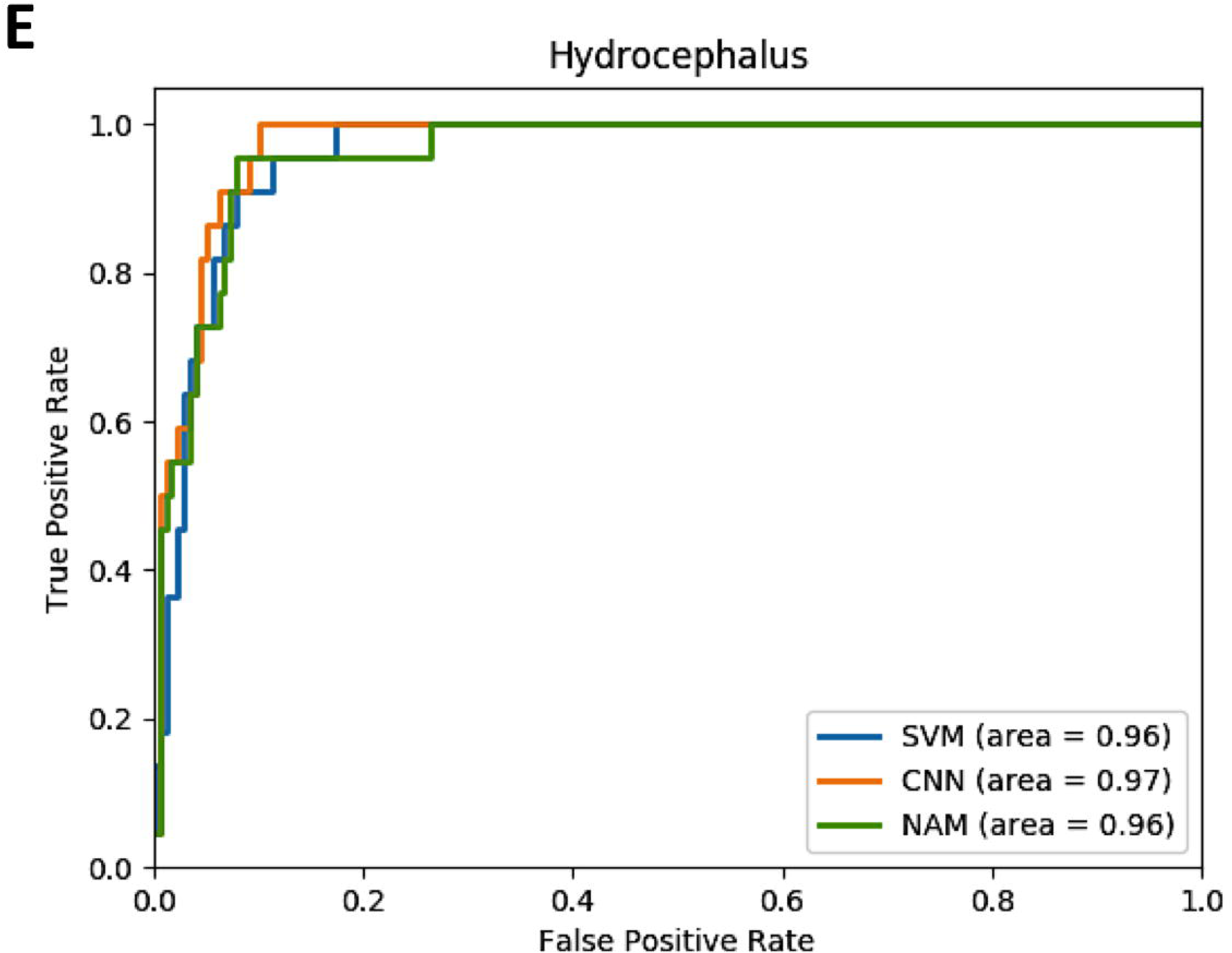
Performance of Machine Learning Algorithms. ROCs of three algorithms for classification of acute and communicable findings; (A) Severity of Study, (B) Acute Blood, (C) Mass Effect, (D) Acute Hydrocephalus, and (E) Acute Stroke. AUC values are denoted by “ area=”. ROC, receiver operator curves; AUC, area under curve.

## Discussion

The purpose of this study was to evaluate the performance of both non-neural and neural network based NLP methods on the document-level extraction of acute and communicable findings in a sample of ICU head CT reports without linkage to any medical language ontologies. The results show that neural network models (CNN and NAM) tend to generally outperform the comparison non-neural models (SVM) for all five classes. The accuracies achieved by neural network models in the five classes for identification of acute and communicable findings range from 0.90 to 0.94. AUCs of all three methods for all classes were above 0.92 indicating excellent performance over multiple sensitivities.

Previous studies have focused on rules-based NLP with variable linkage to established medical ontologies, with accuracies ranging from 80-90% depending on type of radiology report evaluated (e.g. knee MRI or chest radiographs).^3-7, 16^ Non-neural machine learning based NLP methods in radiology have adapted *n*-gram modeling,^17^ naïve Bayes classification,^18^ and, more recently, support vector machines (SVM),^1, 10^ and bag-of-words representation for classification.^19^

However, because some of these published methods have been dependent on mapping to existing medical language ontologies^6, 7^ such as Systematized Nomenclature of Medicine – Clinical Terms (SNOMED-CT),^20^ RadLex for radiology specific lexicon,^21^ or the Unified Medical Language System (UMLS) Metathesaurus,^22^ they have limited use on reports containing language or terms not recognized based on the ontology. They did not have the ability to iteratively “ learn” new variations of terms that describe a finding, recommendation, or desired concept.^16^ For example, there are many ways to say “acute intracranial hemorrhage”; current basic classifier and extraction systems are limited in their ability to recognize any new many variations of these words apart from what is already programmed in the software by humans.

More sophisticated methods such as neural network based deep learning techniques (e.g. convolutional neural networks) have been considered more powerful, able to perform document level classification, and can iteratively learn to improve accuracy,^23, 24^ yet, their performance have not yet been evaluated on radiology reports.

Our results show that the methods used, especially neural network methods have the ability to classify important findings in the head CT report without any need for negation (e.g. differentiating “no stroke” vs. “ there is stroke), linkage to medical ontologies, or word-by-word annotation. Additionally, the neural network models were initially “trained” to evaluate semantic and syntactic patterns on a large un-annotated set of reports (80,000 reports), which is a feature not possible with non-neural machine learning methods like SVMs.

This study is an example of how radiology NLP can be applied to unstructured data (i.e. the radiology report) to extract meaningful information to develop discrete data groups: 1) cohort building for epidemiologic studies, 2) quality assessment for radiology practice, and 3) clinical support services^2^ from clinical databases.

One large group of such clinical databases is electronic health record (EHR) systems. EHRs are replete with large volumes of unstructured data that can be mined for useful population and patient level information.^25^ With increased mandates by federal regulators to demonstrate quality, improve outcomes, and reduce costs,^26^ there is an increasing need to develop scalable and reliable methods of unstructured data mining. Additionally, the Precision Medicine Initiative (PMI)^27^ has spearheaded the need for powerful text mining techniques to promote more nuanced phenotyping of patients and patient populations.^28^

Our study does have some limitations. Although the accuracies and AUCs of the machine learning methods were relatively high, they were not perfect. We did not validate the algorithms on head CT reports from other institutions. We had a relatively modest sample size of 1400 annotated head CT reports. However, human annotation of such reports requires expertise in head imaging and can be laborious. Lastly, our dataset was from a large quaternary hospital’s ICU population. Therefore, we cannot, as yet, verify reproducibility of the algorithms on head CT reports form smaller, community hospitals. The advent of multi-institutional annotated reference sets will likely obviate these limitations.

## Conclusion

We have reported the excellent performance of non-neural and neural machine learning NLP algorithms for the classification of acute and communicable findings on head CT reports from an ICU population. This study’s results show that modern machine learning methods, especially those with neural networks, can help extract meaningful information from unstructured text that is contained in the data warehouses and EHRs. The information discovered by algorithms can be used for outcomes, quality improvement, cost analysis, and operations research.

